# ctSpyderFields: A Python package for visual field reconstruction in spiders

**DOI:** 10.64898/2026.05.28.728173

**Authors:** Massimo De Agrò, Daniele Caradonna, Atal Pande, Egidio Falotico, Lauren Sumner-Rooney

## Abstract

The measurement of visual fields in arachnology has a long-standing history. Given the wide variety of eye positions, orientation and structure, the topic is fundamental for studies of taxonomy, evolution, ecology and behavior. The existing methods for measuring visual fields deploy ophthalmoscopic measurements, which require custom microscopes, anatomical structures like the reflective tapetum, which may not always be present, or the capacity to detect photoreceptor autofluorescence. Here we present the ctSpyderFields python package: a tool for geometrically predicting the visual fields of arachnids from digital images of the lens and retina. The tool uses images coming from computed tomography (CT) scans of specimens, but could be applied to other 3D microscopy techniques, to virtually project the boundaries of the retina through the geometrically predicted nodal point of the lens, deriving a rough per-eye visual field both in cartesian and spherical coordinates. The extracted data can then be used to calculate likely visual field overlap between eyes and angular spans, which can be compared within or between species. We also provide a use case, reporting the visual field data extracted from a museum specimen of *Philaeus crysops*. We propose that the tool will allow a wider comparative analysis of visual fields across spider species, unlocking the potential for a deeper understanding of visual ecology and evolution.

## 2 Introduction

The visual field, or field of view, of an eye describes the region of space it samples. These fields vary in their size, shape, and orientation, and define the spatial limits of information collected by the eye. In animals with two or more eyes, these fields may overlap between eyes, providing a range of potential benefits including increased sensitivity, stereopsis, and redundancy [1]. Animals with many eyes, such as spiders, may have overlap between multiple different eyes on one side of the body, or between bilateral sides. Although the functional implications of these more complex patterns are not studied in as much depth as binocularity, the resampling of certain areas of visual space may imply the likely location of pertinent stimuli, and/or that the different eyes sample the space for different information. As a result, visual field measurements in spiders have a long history, dating back nearly a century. Homann [2] was the first to use the reflective properties of the tapetum and the natural image formation by the eye to deduce optical parameters such as the field of view, focal length, and receptor spacing. His technique was based on direct optical observation of the retina in living specimens, using the eye’s own optics to visualize the retinal mosaic. Land [3] adapted this standard microscope setup into an ophthalmoscope by adding an extra lens, positioned so that it images the back focal plane of the microscope’s objective onto the eyepiece. Because the back focal plane is conjugate with infinity, this setup ensures that the retinal image is viewed as though it were formed at infinity. This modification captures a sharp, angularly calibrated image of the retina. These methods, however, depend on the presence of a reflecting tapetum, which is not present in all spider eyes. Stowasser et al. [4] further refined this approach by exploiting the autofluorescence of the photoreceptors themselves to image the retina, thus broadening the applicability of ophthalmoscopic techniques to a wider range of invertebrate eyes. However, some limitations remain: this method is best applied to live or fresh tissues and requires a custom-built optical system.

In recent decades, micro-computed tomography (micro-CT) has emerged as a powerful and accessible tool for high-resolution, three-dimensional visualization of animal tissues (see [5] for an overview). It has applications in many areas of biology and is increasingly available within academic institutions. Volumetric data from micro-CT can be re-sliced and examined from multiple angles, offering insights into the full anatomical context of the structures. Finally, micro-CT scanning does not require live specimens and is non-invasive, extending its utility to preserved and historically or taxonomically important material. Within vision research, the fundamental biological and optical links between eye structure and function are relatively well understood [6], [7], [8], and inferences about both function and ecology can be cautiously drawn from morphology. As a result, micro-CT methods are already commonly deployed in animal vision research, most notably in the three-dimensional (3D) study of compound eye structure and its functional implications [9], [10], [11].

Here we present a novel procedure to virtually estimate the possible visual field of spiders starting from computed tomography of specimens’ cephalothorax and exploiting known relationships between lens geometry and function. The ctSpyderFields python3 [12] package [13] uses user-segmented retinas and lenses to infer the visual field span and possible overlaps between eyes, also providing convenient data output and visualization tools. Moreover, the package is distributed as fully open source.

## 3 Materials and methods

A full usage guide is provided in the GitHub repository, together with sample code and data [13]. The package employs functions and data structure from other python packages, namely opencv [14], trimesh [15], pandas [16], tqdm [17], matplotlib [18], plotly [19], scipy [20], pyYAML [21], scikit-learn [22], alphashape [23], numpy [24], [25]. This paper instead more generally describes the software usage and the computational solutions employed, together with the description of a sample jumping spider specimen analyzed for the sake of validation.

### 3.1 Specimens’ selection and preparation

To use the python package, first it is necessary to acquire CT scans of spider specimens, and to segment the relevant region of interest (corneal lenses and retinae). For the development of the software, we scanned a variety of spider specimens of the families Salticidae, Lycosidae, and Thomisidae at the Museum für Naturkunde Berlin. This spread was selected to ensure enough variety of lenses and retinas positions and dimensions to ensure generatability of the software, and provides comparability to existing ophthalmoscopic studies of their visual fields [26], [27], [28]. The specimens were stained in 1% iodine by weight in 70% ethanol for three days. Specimens were scanned using a Phoenix nanotom X-ray|s tube (Waygate Technologies, Wunstorf, Germany; RRID:SCR_022582). The cone beam reconstruction was performed using the datos|x 2.2 reconstruction software (Waygate Technologies, Wunstorf, Germany) and the data were visualized in VG Studio Max 3.5 (Volume Graphics GmbH, Heidelberg Germany). The resulting image stacks were imported into Dragonfly version 2022.2.0.1409 [29]. All four cuticular lenses as well as retinae on one side of the head were segmented using the ROI painter tool. Additionally, seven points on the cephalothorax were segmented as orientation markers (used to orient the spiders during the analysis) - anterior to the eyes, posterior to the brain, the dorsal and ventral sides, center of the spider, left of the spider, and right of the spider. These last three markers were only roughly positioned, as their role is to only determine orientation in the 3D reconstruction. Each of the lenses, retinas, and markers were segmented as separate labels, and exported as binary tiff stacks (white voxels considered in the ROI, black voxels as the background).

### 3.2 Loading the segmented areas

Once the binary tiff stack has been acquired, the images can be imported into the software. The CTspyderField package is fully based around the provided “*Spider*” class, which takes in the raw data, performs the computation and eventually outputs the results. The Spider class is itself composed of four object copies of the “Eye” class, which perform the eye-specific computations. Upon first creation of the Spider object, the tiff stack is imported into python using the opencv package [14], converting the images into numpy [24], [25] value matrices (of shape x, y, z, where x and y are the tiff resolution and z is the total number of images). Thereafter, the three-dimensional coordinates of every voxel with a value > 0 (i.e., white in the image) is recorded, together with a user-provided voxel size, as recorded from the ct-scan itself (this will be used to translate the computed measures into real-world units). By the end of this process, the package will have collected for each eye two lists of three-dimensional coordinates, one for the lens and one for the retina, plus one list of seven coordinates, one for each body marker. For ease of computation and memory efficiency, these lists of coordinates are converted in “*PointCloud*” objects from the package trimesh [15].

### 3.3 Orienting the specimen

To ensure a common reference frame for the visual field projection, the *PointClouds* need to be reoriented into a standard relative to the head position, rather than the CT-scan default one, as of course specimens are mounted in the scanner without a universal positioning. To do so, we employ the seven cephalothorax markers. To start, the front-back markers are aligned with the x-axis, with the front towards +x and the back towards -x. Then, the z axis is derived as coplanar of the bottom-top marker-pair, but fully perpendicular to the x-axis. Then, the y-axis is derived as the perpendicular axis between x and z. This set of rotations makes so that the spider is facing +x, with the top of the cephalothorax pointing towards +z and its right side pointing towards +y. Lastly, the full system is translated such that the center marker is set as the cartesian origin (0,0,0). At this stage, the package saves the resulting roto-translational matrix, that will then be applied at the end of computation in order to visualize the results in the common reference frame.

### 3.4 Identification of the lens nodal point

As a key step of the computation, we need to identify the nodal point of the corneal lens. For corneal lenses with a uniform refractive index, or with concentric layers of refractive indices, this can be approximated at the center of curvature of the lens surface [30]. To find this, we fit an imaginary sphere to the outer surface of the lens. This first required us to identify and isolate only the points on the outer surface of the lens points cloud, which should be used to compute the sphere, and discard points on the proximal side of the lens.

To do this, we used the position of the retina. It is always the case that the outer surface of the lens is that furthest from its associated retina. Therefore, by employing the vector connecting the retina and lens centers, we can use the plane perpendicular to it to bisect the lens surface into proximal (i.e., half near the retina) and distal (i.e., half far from the retina) of the cephalothorax. From here, we fitted a sphere to the outside lens surface, using the least square methods using the code provided in [31], and use its center as the nodal point. In some cases, where for example the lens surface is particularly flat, the sphere-fitting algorithm may not converge, or find spheres with near-infinite radius. In such cases, the focusing power provided by the lens is presumed to be minimal and any further use of the ctSpyderFields package should be extremely cautious. The user can manually input a ‘dummy’ nodal point as a spot on the retina-lens axis at a manually defined distance from the lens surface. This will allow the analysis to run and output plausible centroids for the visual field of each eye; however, all outputs concerning visual field spans and overlaps will be overestimates. Setting the dummy nodal point as being immediately proximal to the lens will reduce, but not mitigate, this effect, and does not reflect what is optically occurring in the eye.

Once all lens nodal points have been found, the roto-translation derived in step 2.3 is applied to the whole Spider object.

### 3.5 Retinal projection

Finally, the visual field can be estimated by iterating through every point in each retina cloud, and project a line crossing the nodal point of the respective lens. The logic behind this is that these lines should intersect the outer lens surface at an angle of 90°, and incoming light at this angle is not refracted. In theory, these lines could go towards infinity, but for computational efficiency they are projected until hitting an imaginary sphere with a radius of 150 mm. This value is chosen by default, but it can be specified by the user and it should be selected according to the species estimated depth of field. The dimension of the sphere does not influence the results regardless, as all visual field characteristics are expressed either in spherical coordinates or as percentages of the full ideal visual field. Multi eye overlap instead may be influenced by the choice of this value, as a longer projection will naturally result in a bigger proportion of the visual fields overlapping each other. The intersection between the lines and the sphere are computed using a ray-tracing algorithm from [32], resulting in a set of 3D coordinates lying on the sphere surface. Thereafter, these points are converted into spherical coordinates (radius, azimuth and elevation). Azimuth (-π, π) and elevation (-π/2, π/2) instead represent the elective system of reference for describing visual field characteristics.

### 3.6 Visual field contours

At this stage, we have projected every original point in the retina cloud. However, to find the spatial limits of the visual field, we need to isolate the perimeter of this cloud. To achieve this, we apply the alpha-shape algorithm from the alphashape python package[23]: a generalization of the convex hull algorithm also capable of describing concave contours. Applying the alpha-shape [33] algorithm on spherical coordinates allow us to treat the surfaces as if they were flat, simplifying the calculation greatly in respect to 3D. This provides and plots the visual fields for one side of the spider, which can then be mirrored over the xz-plane for acquiring the contralateral visual fields. The package can output in csv format both the list of contour points of each visual field in spherical coordinates, along with summary data including vertical and horizontal span for each eye, total coverage, and orientation of each visual field. The Spider object can also be saved as a pickle file for future analysis without having to re-extract point clouds from the images.

## 4 Results

As a proof of concept, we segmented and analyzed the visual fields of a jumping spider, *Philaeus chrysops* (ZMB 19219), taken among the samples scanned for the initial software design. Jumping spiders have received significant research attention due to their outstanding visual capabilities and behaviors, and their visual fields have been measured via ophthalmoscopy in multiple species [34], [35], [36]. In salticids, the largest pair of eyes is the forward-facing anterior median eyes (AMEs). The retinas are cup-shaped and are positioned at the end of a long, movable eye-tube. They project a narrow, boomerang-shaped visual field, as already described via ophthalmoscopy [3], [37]. This represents a challenging and unusual structure, which tests the capabilities of the ctSpyderField package. The posterior lateral eyes (PLEs) are the smallest pair, considered to be vestigial in many species. The anterior lateral (ALEs) and posterior median eyes (PLEs) are similarly sized, and they respectively project their visual fields towards the front and towards the back of the cephalothorax [27]. This pattern is consistent across a variety of species, and has been behaviorally validated [38], [39], [40], providing a robust benchmark for our method.

The *Philaeus chrysops* specimen was scanned at a voxel size of 0.003 mm. We segmented eyes and retinas from the right side only, and then mirrored for acquiring information about binocular fields. The data was then extracted from dragonfly and loaded into the python environment. As measured by the distance between front and back, left and right, and top and bottom markers respectively, the *P. chrysops* carapace was 3.49 mm long, 2.76 mm wide and 2.95 mm thick.

### 4.1 Lens and retina measurements

The calculated AME lens volume was 0.0914 mm^3^. The sphere fitted to the lens surface had a radius of 0.3844 mm. The ALE lens volume was 0.0166 mm^3^, and the fitted sphere radius was 0.2127 mm.

The PME lens volume was 0.00026 mm^3^, while the fitted sphere radius was 0.0754 mm. The last lens, the PLEs, had a volume of 0.0108 mm^3^, and the fitted sphere radius was 0.1862 mm.

**Figure 1.**
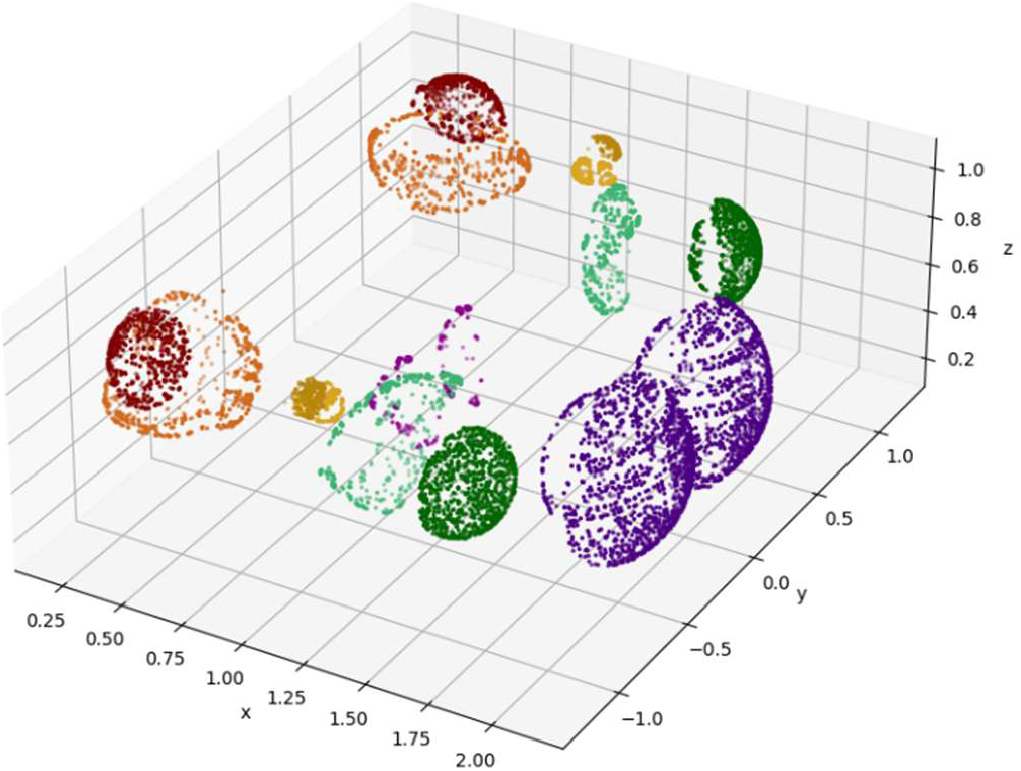
3D plot produced using the dot coordinates of both lenses and retinas of the Philaeus chrysops eyes. In purple, AMEs, In green, ALEs, in yellow, PMEs, in red, PLEs. The cephalothorax is oriented according to the standard, and as such the front of the spider is towards +x, while the top is towards +z. The anatomical data has been collected for the right-side eyes, while the left side here depicted is a mirror image.

As stated above, in Salticidae the large AMEs lenses are coupled with small, cup-shaped retinas with four layers at the back of the long eye-tubes [41]. This resulted in a volume of 0.0011 mm^3^. ALEs and PLEs are instead larger and more classically structured in respect to other spider families. The retina volume was respectively 0.0069 mm^3^ and 0.0206 mm^3^. The PME retina was smaller, with a volume of 0.0009 mm^3^.

### 4.2 Visual field projections

In line with existing data [27], the AME visual fields are the smallest, and are boomerang shaped. The total span of each eye field is 0.4722 radians of elevation and 0.2159 radians of azimuth. Within this, the ‘tallest’ point measured 0.312 radians in elevation, observed at 0.096-0.114 radians azimuth, at the center of the visual field. In terms of azimuth, the maximal span is 0.1 radians, observed at an elevation between 0.254 and 0.271 radians, at the very top of the field. The AME field has its centroid at 0.1045 azimuth and 0.05 elevation.

There was no binocular overlap between the two AME fields. However, the retinas of these eyes are movable, which could generate some overlap between their fields during active vision [42], [43]. The AME visual fields are fully contained within the ALEs fields, with a 100% overlap, but there is no overlap between the AME fields and the visual fields of either posterior eye pair. With respect to the complete spherical field around the spider’s cephalothorax, one AME field occupies 0.16%.

**Figure 2.**
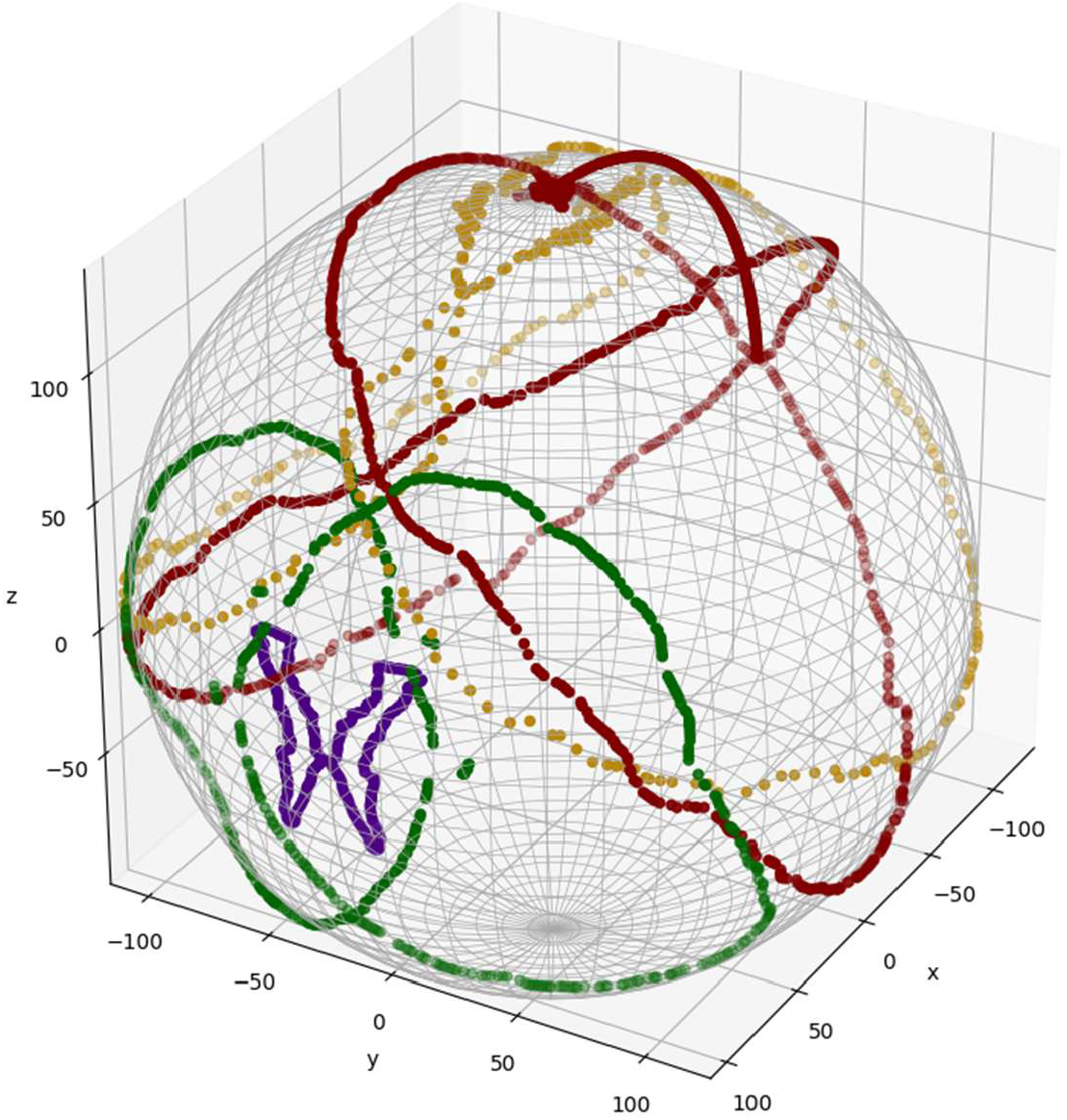
Visual fields projected onto the 150mm radius sphere. The spider is virtually facing towards +x. In purple, AME fields; in green, ALE fields; in yellow, PME fields; in red, PLE fields. Note the boomerang-shaped visual field of the AMEs, and the centrally-overlapping fields of the ALEs.

**Figure 3.**
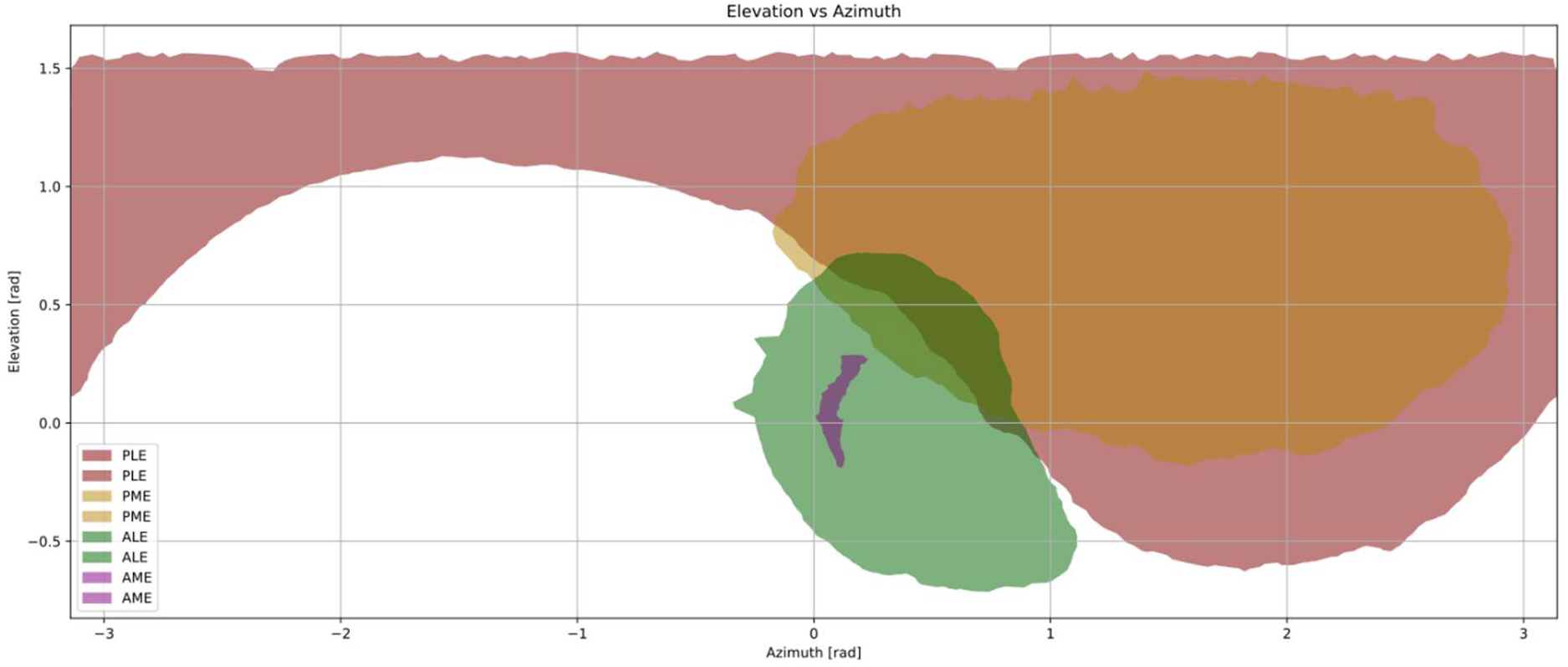
Visual field in spherical coordinate system. Only the right side is shown here for visual clarity. Note how the PLE field spans across the whole azimuth range above 1.2rad.

The ALE visual fields are forward-facing and drop-shaped. The total span of the right field ranged from -0.34 to 1.11 radians in terms of azimuth (total 1.45 radians), and from -0.71 to 0.72 radians in terms of elevation (total 1.43 radians). The maximal ALE elevation was 1.358 radians, observed at 0.45-0.46 radians azimuth. The maximal azimuth width was 1.168 radians, observed at 0.06-0.08 radians elevation. The ALE field has its centroid at 0.4004 azimuth and -0.035 elevation.

The ALE fields overlapped frontally to the spider for 26.8% of their total area. Moreover, they overlapped 22.3% with the ipsilateral PME field, and 15.3% with the ipsilateral PLE. Each ALE covered 6.8% of the complete spherical field, while the combined ALEs field covered 11.9%. The PME fields were large, dorsal-facing and generally circular. The total elevation span was -0.18 to 1.48 radians (total 1.66 radians), and the total azimuth span was -0.17-2.95 radians (total 3.12 radians), and from. The maximum elevation was 1.63 radians, observed at an azimuth between 1.53 and 1.55 radians, and the maximum azimuth width was 3.106 radians, observed at an elevation between 0.76 and 0.78 radians, i.e., the field was tallest and widest roughly at its center. The PME field has its centroid at 1.5327 azimuth and 0.9332 elevation. The PMEs had a very small binocular overlap, amounting to only 2.8% of their total area. Moreover, each PME field overlapped 7.8% with the ipsilateral ALE field and 19.5% with the ipsilateral PLE. The one PME field covered 19.5% of the full spherical field, while their combined fields covered 38.5%.

The PLE fields are the largest of all, spanning from the anterior to posterior of the spherical field around the spider. This wide coverage of the PLEs makes it difficult to properly describe the field in terms of azimuth and elevation, as it crosses the poles of the sphere and the 180-degree mark in terms of azimuth. For this reason, the total field spanned almost the full spherical area around the spider, spanning from -3.142 to 3.142 radians in terms of azimuth (total 6.284 radians), and from -0.62 to 1.571 radians in terms of elevation (total 2.19 radians).

**Figure 4.**
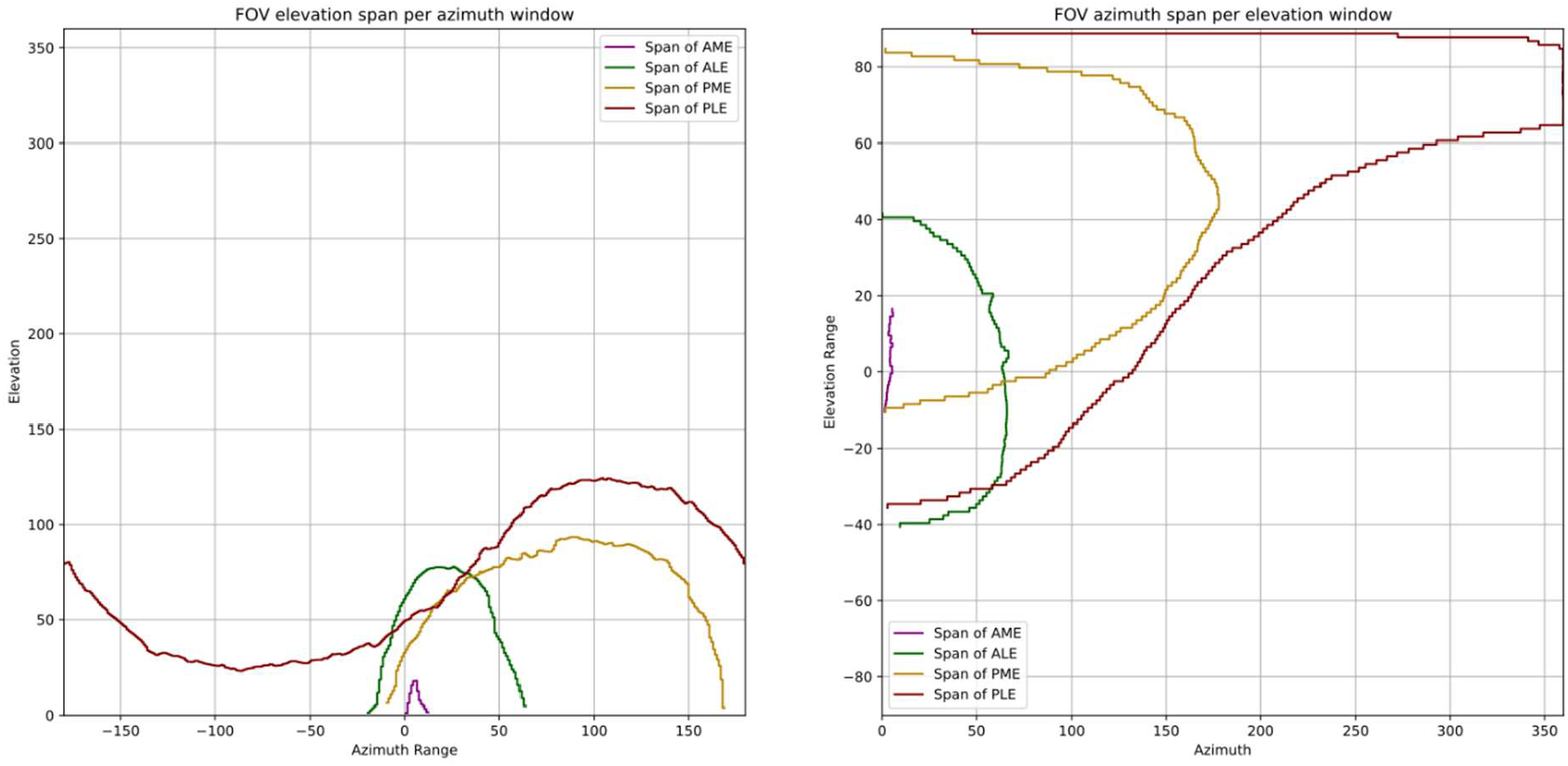
Plot of angular span of each visual field across elevation and azimuth steps, in degrees. The left plot shows the elevation spans across each azimuth range. Note for example that the maximal elevation span of the AME can be appreciated at an azimuth of 5-6 degrees in respect to the exact spider front. The right plot shows the azimuth span over each elevation range. Note for example the full 360°azimuth span of the PLEs between around 65 and 85 degrees.

The PLE maximal azimuth width was 6.284 radians, observed at 1.22-1.23 radians elevation, i.e., the PLE field formed a near-circle near the dorsalmost point of the sphere. The maximal elevation was 2.171 radians, observed at 1.81-1.83 radians azimuth. Here the effect was the opposite, as the fields are intersected by the pi/2 elevation line that marks the bounds of our spherical coordinate system. The field at this azimuth continues on the opposite side of the sphere, so between -1.81 and -1.83 radians, and has an elevation span of 0.466 radians. As such, the maximal elevation span of the PLEs is in fact 2.637 radians. The PLE field has its centroid at 2.8225 azimuth and 1.088 elevation. The PLEs had a large binocular overlap, amounting to 53.15% of each other. This is almost fully located at the dorsal extreme of the spherical field. They also overlap considerably with PMEs, for 49.65% of their full surface, and for a small section with the ALEs, specifically 2.76%. A single PLEs covers 38.01% of the full spherical field, while the binocular field covering a total of 55.82%.

## 5 Discussion

Here, we presented the ctSpyderFields package, an open-source tool for the virtual projection of visual fields of spiders as derived from ct-scanning. We also presented a use case of the software, describing the visual field of the jumping spider *Philaeus chrysops*.

### 5.1 Advantages

The ctSpyderFields package offers significant advantages over traditional methods of visual field estimation commonly used in spider research. As stated in the introduction, ophthalmoscopic methods require specialized equipment and delicate components. They may not work universally on all spiders, depending on the presence of a tapetum or sufficient autofluorescence. Behavioral procedures are even more limited, as they require a sufficiently consistent behavioral output (like the pivots towards moving objects in jumping spiders, see [44], [45]) while still providing a very low spatial resolution in the output data, often limited to the maximal horizontal span [39], [40]. Here, we could estimate visual fields from collection specimens, without any need to capture live spiders. This massively increases the reach of the tool, as across Europe the variety of species already acquired and available as sample could allow for an almost universal survey of visual fields across the whole Order. The procedure is also quite tolerant to the sample quality: some or our specimens has suffered significant dehydration damage, but the retina and lenses were conserved enough that allowed for anatomical reconstruction, which sufficed for the application of the procedure.

Other than the wide applicability, the package also offers an unprecedented level of detail and comparability between specimens. This is because the procedure fully gets away with the need of human intervention on the positioning of the animal in the specialized machinery, and take cares of scaling and orientation fully virtually. The level of precision achievable at such a small scale (given the average size of the spiders) is massively increased by the digital approach. The data outputted is also of a much finer detail that what could be achieved with traditional methods, based on the voxel size of the scan. On this note, given the fact that the system starts computation from segmented tiff stacks, the same procedure could be applied starting from any other software or type of volumetric anatomical approach, with possibly also increasing the available resolution (e.g., FIB-SEM).

### 5.2 Disadvantages

The package is, of course, not free from disadvantages and limitations. On a practical level, the first step of the procedure still relies on human input for eye and retina segmentation. This introduces possible sources of noise, as manual procedures are vulnerable to human error, and increases the time required to collect the data. This in turns limits the scale of applicability: scanning and segmenting hundreds of specimens will take a significant human effort compared to the few seconds or minutes needed to perform the virtual computations. However, the rapid development of automatic or AI-assisted segmentation tools may cut greatly the working time-per-specimen in future [46].

In order to maintain generalizability, the package is also required to make some general assumptions. First, our geometric approach uses the center of curvature of the lens as its nodal. In some cases, where lens curvature is very shallow, this point may fall behind the retina. Application of the package to these underfocussed eyes will fail to truly represent the spider’s visual field. Second, the position and size of the retina may be affected by specimen preservation methods. We selected a scanned specimen that exhibited minimal dehydration, from inspection of the internal tissues, but alcohol can cause significant tissue shrinkage that may distort the retina. Third, as mentioned above, the retinas of the AMEs are mobile and have up to several dedicated muscles; the orientation of their visual fields are therefore also mobile. Finally, we have used the nodal point projection to dramatically simplify the problem of predicting the visual field, as we are hypothetically restricting our estimates to incoming light perpendicular to the lens surface; thus, refraction is not accounted for. In a focusing lens, refraction increases the field span, so this renders our estimates conservative. These are considerable compromises, but we hope that it may be used as a comparative tool that is perhaps more accessible than high-end ophthalmoscopy.

### 5.3 Conclusions

Overall, the ctSpyderFields package is an open-source, accessible and user-friendly tool for the identification of visual fields of spiders. The system is robust enough to offer comparability between specimens and techniques, while still maintaining enough flexibility for being useful in edge cases. We believe that the adoption of such a tool will unlock a level of visual field comparison between phylogenetically far and near species, allowing for the discovery of new patterns of eye evolution in this Order.

## 6 Acknowledgments

The development of this software was possible due to a Synthesys+ Transnational Access Grant to MDA. LSR and AP are supported by the DFG Emmy Noether Programme (SU 1336/1-1 and SU 1336/1-2).

